# Carbonyl Reductase 1: a novel regulator of blood pressure in Down Syndrome

**DOI:** 10.1101/2024.05.17.594787

**Authors:** Alexandra J Malbon, Alicja Czopek, Andrew M Beekman, Zoë R Goddard, Aileen Boyle, Jessica R Ivy, Kevin Stewart, Scott Denham, Joanna P Simpson, Natalie Z Homer, Brian R Walker, Neeraj Dhaun, Matthew A Bailey, Ruth A Morgan

## Abstract

**Background:** Approximately one in every 800 children is born with the severe aneuploid condition of Down Syndrome (DS), a trisomy of chromosome 21. Low blood pressure (hypotension) is a common condition associated with DS and can have a significant impact on exercise tolerance and quality of life. Little is known about the factors driving this hypotensive phenotype and therefore therapeutic interventions are limited. Carbonyl reductase 1 (CBR1) is an enzyme contributing to the metabolism of prostaglandins, glucocorticoids, reactive oxygen species and neurotransmitters, encoded by a gene (*CBR1*) positioned on chromosome 21 with potential to impact blood pressure.

**Methods:** Utilising genetically modified mice and telemetric blood pressure measurement, we tested the hypothesis that CBR1 influences blood pressure and that its overexpression contributes to hypotension in DS.

**Results:** In a mouse model of DS (Ts65Dn), which exhibit hypotension, CBR1 activity was increased and pharmacological inhibition of CBR1 increased blood pressure. Mice heterozygous null for *Cbr1* had reduced CBR1 enzyme activity and elevated blood pressure. Further experiments indicate that the underlying mechanisms include alterations in sympathetic tone and prostaglandin metabolism.

**Conclusions:** We conclude that CBR1 activity contributes to blood pressure homeostasis and inhibition of CBR1 may present a novel therapeutic opportunity to correct symptomatic hypotension in DS.

## Introduction

Down Syndrome (DS) is the most common chromosomal disorder, affecting 1 in every 792 babies born^1^. 95% of people with DS have a trisomy of chromosome 21 with resultant effects on development. Patients with DS are at risk of comorbidities including hypothyroidism, sleep apnoea, obesity, metabolic syndrome, psychiatric disorders and Alzheimer’s disease^2,3^. Low blood pressure – hypotension - is common in both children and adults with DS^4–6^. This hypotension results in lower cardiorespiratory fitness and an inadequate blood pressure response to sub-maximal and maximal exercise^7,8^ limiting the ability to participate in many activities^9^ which in turn impacts quality of life. DS patients also commonly have non-dipping nocturnal blood pressure and heart rate which may contribute to sleep disorders and an increased risk of cardiovascular events^10–12^. There is also a well-documented association between low blood pressure and the development of Alzheimer’s disease which is particularly common in patients with DS^5^. Baseline hypotension also makes the interpretation of blood pressure as a diagnostic tool for detecting other co-morbidities challenging^4^. Despite these impacts, the pathogenesis of hypotension in DS has not been elucidated; some have suggested that it is due to autonomic dysfunction since clinical studies report reduced sympathetic and increased parasympathetic tone in patients with DS^13–17^.

*CBR1,* the gene encoding the ubiquitously expressed enzyme carbonyl reductase 1^18^, is located in the ‘Down Syndrome critical region’ of chromosome 21, the region that co-segregates with many of the developmental features of DS^19,20^. CBR1 is a complex enzyme with a number of substrates and is most often studied for its role in metabolism of therapeutics such as doxorubicin^21^. CBR1 plays a critical role in cellular homeostasis and blood flow regulation by preventing the accumulation of reactive oxygen species, vasoconstrictor prostaglandin E_2_ and neuroactive metabolites such as monoamine oxidase inhibitor and endogenous indoles^22–25^. Recent data suggest that CBR1 activity is important in regulating renal blood flow via prostaglandin metabolism^26^. Our work has also shown the role of CBR1 in tissue metabolism of glucocorticoids^27^ and its impact on glucose homeostasis in lean mice^28^. In this study we used a transgenic murine model of *Cbr1* deletion, as well as pharmacological inhibition of CBR1 in a murine model of Down Syndrome, to address the hypothesis that CBR1/*Cbr1* plays a role in blood pressure regulation and that dysregulation of CBR1 contributes to hypotension in DS. We also explore the potential mechanisms by which this might occur.

## Materials and Methods

### Animals

Male B6EiC3Sn.BLiA-Ts(1716)65Dn/DnJ (Ts65Dn mice), a model of DS, with littermate controls were obtained from The Jackson laboratory (Stock #005252)^29^. This line contains a partial trisomy encompassing most of the human chromosome 21 orthologous region of mouse chromosome 16^30^, including *Cbr1*^31^. These animals have been well characterised with regards cerebellar volume which is reduced, as in DS^29^. They demonstrate increased locomotor activity and energy expenditure^32^ (33), have reduced blood pressure^33^ and impaired conscious respiration associated with a decreased neural drive^34^. Mice heterozygous for *Cbr1 (Cbr1^+/-^)* were generated as previously described^28^, homozygosity of this gene deletion is foetal lethal^35^. Data from our group has previously shown that this model has an approximately 50% reduction in CBR1 expression and activity^28^. All experiments were performed in accordance with the UK’s Animals (Scientific Procedures) Act under a UK Home Office Project Licence in accordance with EU Directive 2010/63/EU. Mice were maintained according to institutional guidelines, group housed at 21 ± 1 °C; humidity at 50 ± 10 % with a 12-hour light-dark cycle (light period 07:00-19:00) unless otherwise stated. Unless otherwise specified, mice were killed by cervical dislocation. Mice were fed on a diet containing 0.3 % Na and 0.7 % K by weight (RM1 diet, Special Diet Services, United Kingdom) throughout the experiment unless otherwise stated. Blood pressure was measured by telemetry in *Cbr1^+/-^*and *Cbr1^+/+^* littermate controls and in Ts65Dn mice and their littermate controls at baseline and during treatment with hydroxy-PP-Me, an inhibitor of CBR1^36,37^. Synthesis and purification of hydroxy-PP-Me is described in the supplementary materials. Renal function, vascular function, plasma renin, angiotensin and aldosterone were measured in mice heterozygous for *Cbr1* and their wild-type littermate controls (see supplementary methods and results).

### Blood pressure measurement

Ten-week-old male mice (*Cbr1^+/-^, Cbr1^+/+^*, Ts65Dn mice and wild-type littermates (n=8/group)) had PA-C10 radio-telemetry devices (Data Science International, USA) implanted into the carotid artery under isoflurane anaesthetic (4 % induction, 2-3 % maintenance). Buprenorphine (0.1 mg/kg Vetergesic; Ceva Animal Health Ltd, Libourne, France) was administered subcutaneously prior to recovery and per os (vetergesic jelly) for the first four days. Mice underwent a one-week post-surgical recovery period as basal diurnal rhythmicity of the measures was re-established. Data were obtained for the following 7 days. For the duration of the experiment, five consecutive one minute blood pressure and heart rate readings were taken every 30 min at an acquisition rate of 1kHz. Ts65Dn mice and their wild-type controls then received hydroxy-PP-Me for 1 week during which data were collected. Hydroxy-PP-Me was administered intraperitoneally at a dose of 30mg/kg based on previously published data^37^. Previous work from our group showed there was no effect of intraperitoneal injection alone on blood pressure^38^. *Cbr1^+/-^* mice and *Cbr1^+/+^* littermates did not receive the CBR1 inhibitor but did receive a high-salt diet (3% Na) for 7 days (see supplementary data).

### CBR1 activity

CBR1 activity, as measured by reduction of the substrate doxorubicin, was quantified in hepatic or cardiac cytosol from Ts65DN animals and their littermate controls with or without administration of hydroxy-PP-Me, as previously described^39–41^. Briefly cytosol from homogenised tissue was extracted by ultracentrifugation, the protein quantified by Bradford protein assay. Cytosol was incubated with 50 µM doxorubicin, the reaction was started by addition of co-factor NADPH whose oxidation was measured at 340 nm at 37°C over 3 minutes. Enzymatic velocities were calculated by linear regression of the change in absorbance over time.

### Urine collection and analysis

For collection of urine, mice were housed in metabolic cages for 48 hours. Urinary catecholamines adrenaline and noradrenaline were measured by enzyme linked immunoassay (ELISA) (CatCombi ELISA Kit, Creative Diagnostics, DEIA1663). Prostaglandin E_2_ was measured by ELISA (Cayman Chemical, 514531) in urine from 8-week-old *Cbr1*^+/-^ mice and *Cbr1^+/+^* littermates (n=8/group) according to the manufacturer’s protocol. Urinary 8-hydroxy-2’-deoxyguanosine (8-OHdG) was measured by ELISA (Abcam, ab201734) according to manufacturer’s instructions.

### Markers of oxidative stress

Plasma was collected from a subset of animals at cull. Brains were harvested at post-mortem, snap frozen in liquid nitrogen and stored at −80 °C. Total anti-oxidant capacity was measured in plasma using a colorimetric assay based on reduction of ferric ions (Fe^3+^) to ferrous ions (Fe^2+^) using a phenanthroline substance according to manufacturer’s instructions (ThermoFisher EEA022). Malondialdehyde (MDA) was measured in plasma and brain homogenate by quantifying the adduct generated when MDA in the sample reacts with thiobarbituric acid (TBA) (Abcam, ab118970, Lipid-Peroxidation Kit).

### Plasma analysis

Plasma aldosterone, corticosterone and 11-dehydrocorticosterone were measured by liquid chromatography tandem mass spectrometry as previously described^28^. Plasma renin was measured by ELISA (Abcam, ab193728).

### Statistical Analysis

Power calculations were used to determine sample size (G*Power ^42^) for reliable detection of differences in blood pressure as measured by telemetry. They were based on previously published differences in blood pressure between Ts65Dn mice and their wild-type littermates^33^. A sample size of 7/group was determined to be sufficient to give 80 % power to detect a difference with a significance of P < 0.05 using Cohen’s d effect size; we used 8 animals/group to allow for any complications of telemetry but we did not have to exclude any animals from analysis. For the renal function and tissue analysis we used 6-9 animals/group.

All data were tested for normality using the Kolmogorov-Smirnoff normality test, and the appropriate parametric or nonparametric statistical tests were used accordingly. All statistical tests used were two-tailed. Statistical comparisons were made using a Student’s t-test or Mann-Whitney U test or two-way ANOVA tests with appropriate post hoc tests for multiple groups. The asterisks in the figures indicate statistical significance: \**P* < 0.05, \*\**P* < 0.01, and \*\*\**P* < 0.001. All graphs were plotted with GraphPad Prism software or R ggplot. Blood pressure data were analysed in two ways: first by comparison of the medians of blood pressure and heart rate during the inactive and active periods; and second by cosinor analysis which takes into account the circadian rhythm of these measures. Cosinor analysis was conducted and visualised using the R package Circacompare and Limoryde^43,44^.

## Results

### Blood pressure in *Cbr1^+/-^* mice

Mice heterozygous for *Cbr1* had increased median systolic pressure during both the active and inactive periods and increased diastolic and mean arterial pressure during the inactive phase compared to *Cbr1^+/+^* littermate controls (Table 1). There was no difference in median heart rate between *Cbr1^+/-^* and *Cbr1^+/+^* littermate controls.

**Table 1:** Blood pressure and heart rate of mice heterozygous for *Cbr1* (*Cbr1^+/-^)* and their littermate controls (*Cbr1^+/+^)* during the inactive and active period. Data are median and interquartile range. Genotypes were compared using a Mann-Whitney U test.

|  | Inactive Period |  |  | Active Period |  |  |
| --- | --- | --- | --- | --- | --- | --- |
|  | *Cbr1*<sup>+/+</sup> | *Cbr1*<sup>+/-</sup> | *P-value* | *Cbr1*<sup>+/+</sup> | *Cbr1*<sup>+/-</sup> | *P-value* |
| **Systolic (mmHg)** | 110.2 (105.8, 113.5) | 115.0 (111.9, 116.6) | <0.0001 | 125.1 (122.5, 129.2) | 127.8 (124.3, 131.1) | <0.05 |
| **Diastolic (mmHg)** | 80.31 (76.47, 84.92) | 83.26 (81.42, 85.19) | <0.05 | 94.53 (89.77, 98.67) | 95.97 (94.36, 98.85) | 0.14 |
| **MAP (mmHg)** | 91.17 (86.45, 94.39) | 93.96 (91.96, 95.73) | <0.001 | 105.9 (100.9, 108.4) | 106.6 (104.7, 110.1) | 0.09 |
| **Heart Rate (bpm)** | 455.9 (434.0, 479.5) | 473.5 (442.3, 496.4) | 0.05 | 521.2 (501.8, 553.9) | 530.1 (512.8, 561) | 0.16 |

mmHg – millimetres of mercury; MAP - mean arterial pressure; bpm – beats per minute

The blood pressure and heart rate of both *Cbr1^+/-^* and *Cbr1^+/+^* littermate controls could be modelled with a cosine curve indicating a circadian rhythm, as expected. The rhythm-adjusted mean (MESOR, midline estimating statistic of rhythm) of systolic, diastolic and mean arterial pressure was increased in *Cbr1^+/-^* mice compared with *Cbr1^+/+^* controls (Fig 1, Table 2). There was no difference in the amplitude between the groups for any blood pressure parameter measured, indicating that the increased blood pressure in *Cbr1^+/-^* mice was accompanied by a similar sized inactive phase dip but from a higher active phase pressure than that of *Cbr1^+/+^*controls (Fig 1, Table 2). The rhythm adjusted mean of heart rate was significantly higher in *Cbr1^+/-^* mice compared with *Cbr1^+/+^* controls (Fig 1, Table 2). The amplitude did not differ between the groups for heart rate indicating that the heart rate of *Cbr1^+/-^*mice dipped in the inactive phase but still remained relatively elevated compared with wild type controls (Fig 1, Table 2).

**Fig. 1:**
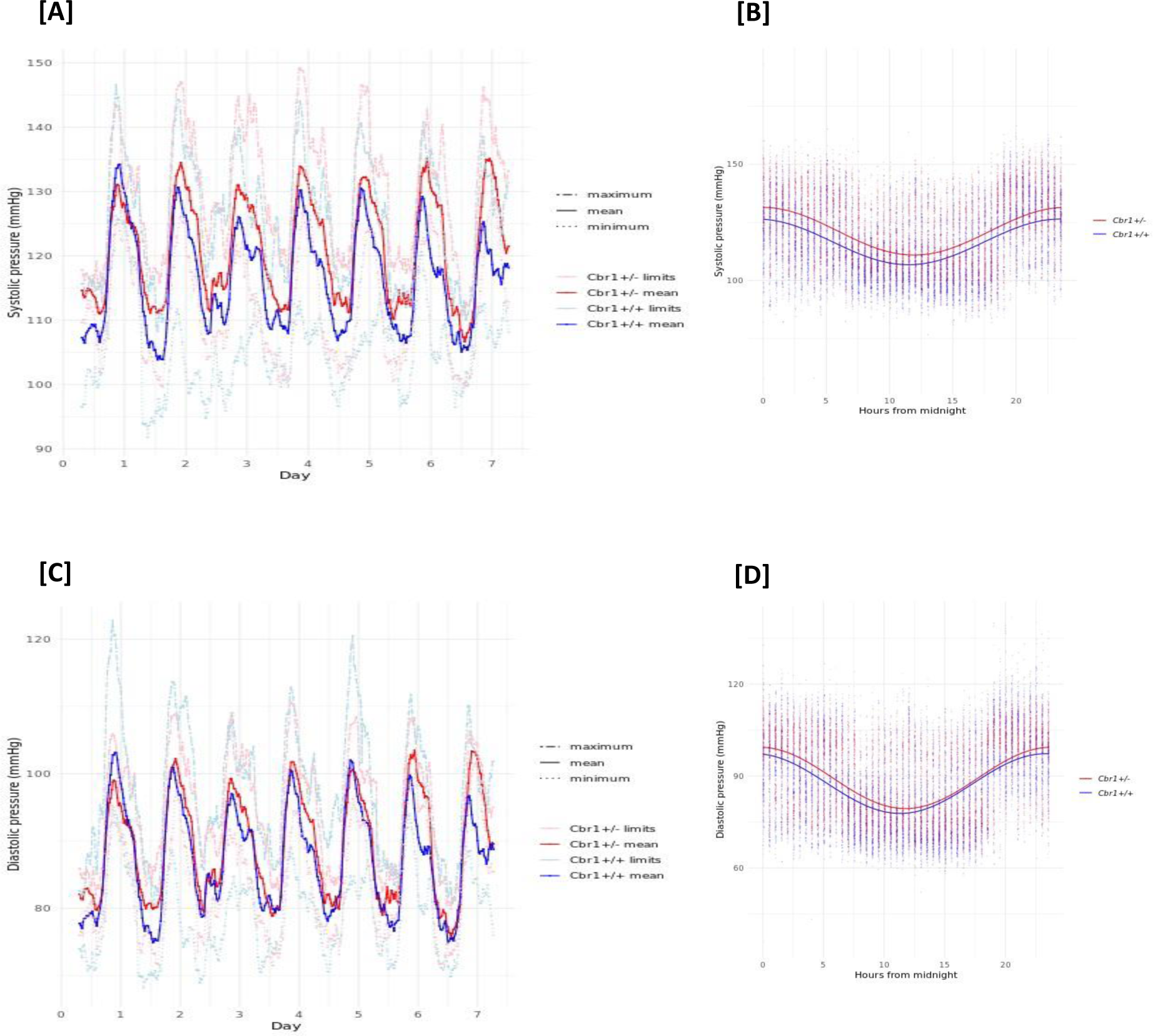
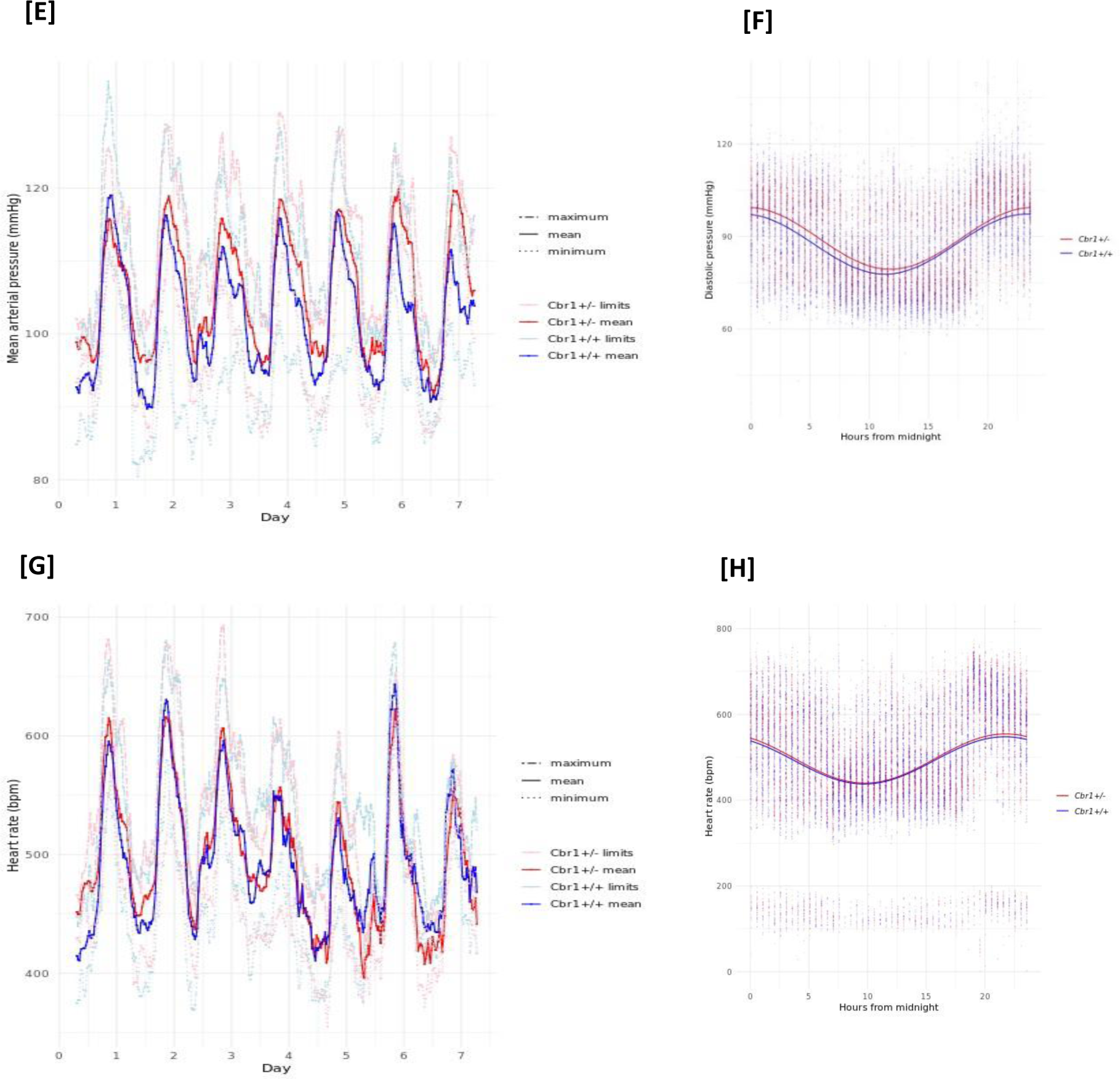
Cbr1 deletion results in elevated blood pressure regardless of cardiac or circadian phase. The left hand column [A, C, E and G] show the five-hour rolling averages and minimum and maximum systolic, diastolic and mean arterial blood pressure, and heart rate of wild type mice (blue) and mice heterozygous for Cbr1 (red) (n=8/group). [B, D F and H] show the cosinor curves and spread of data points for the 7-day measurement period for systolic, diastolic and mean arterial pressure, and heart rate.

**Table 2:**
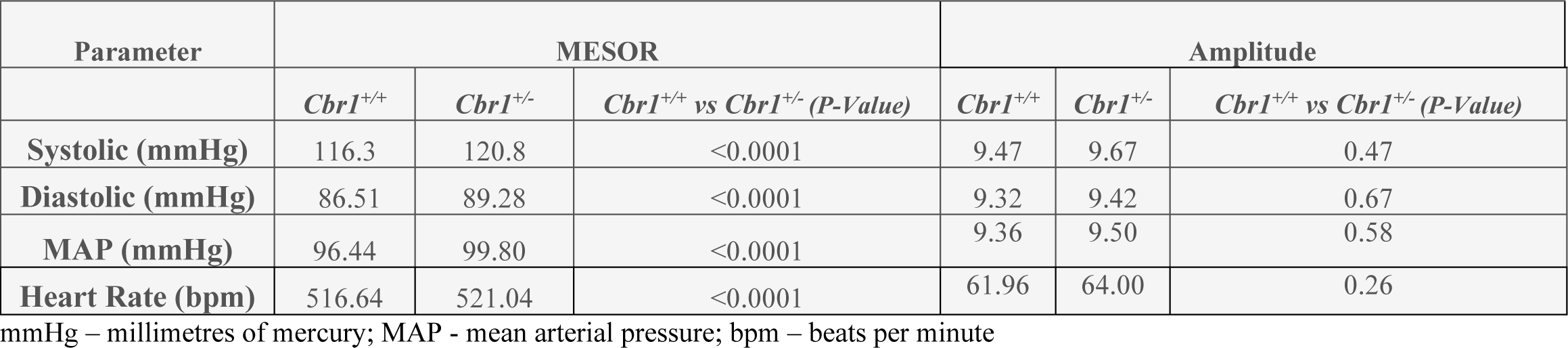
Cosinor analysis of blood pressure and heart rate measured by telemetry in *Cbr1^+/-^* mice and their *Cbr1^+/+^* littermate controls. The rhythm-adjusted mean (MESOR), amplitude of each parameter and the outcome of statistical comparison by Mann-Whitney U test are shown.

### Inhibition of CBR1 in a mouse of model of DS

We hypothesised that a mouse model of DS, Ts65Dn, would have relative hypotension and that pharmacological inhibition of CBR1 would increase blood pressure.

We first confirmed that Ts65Dn mice had higher hepatic and cardiac mRNA levels and CBR1 activity (Fig. S1 and Fig 2) compared with littermate controls. We then determined the extent of inhibition of CBR1 activity by the drug. Administration of the selective CBR1 inhibitor, hydroxy-PP-Me reduced hepatic CBR1 activity in Ts65Dn mice to equivalent to the wild type mice but did not reduce cardiac CBR1 activity (Fig 2).

**Fig. 2:**
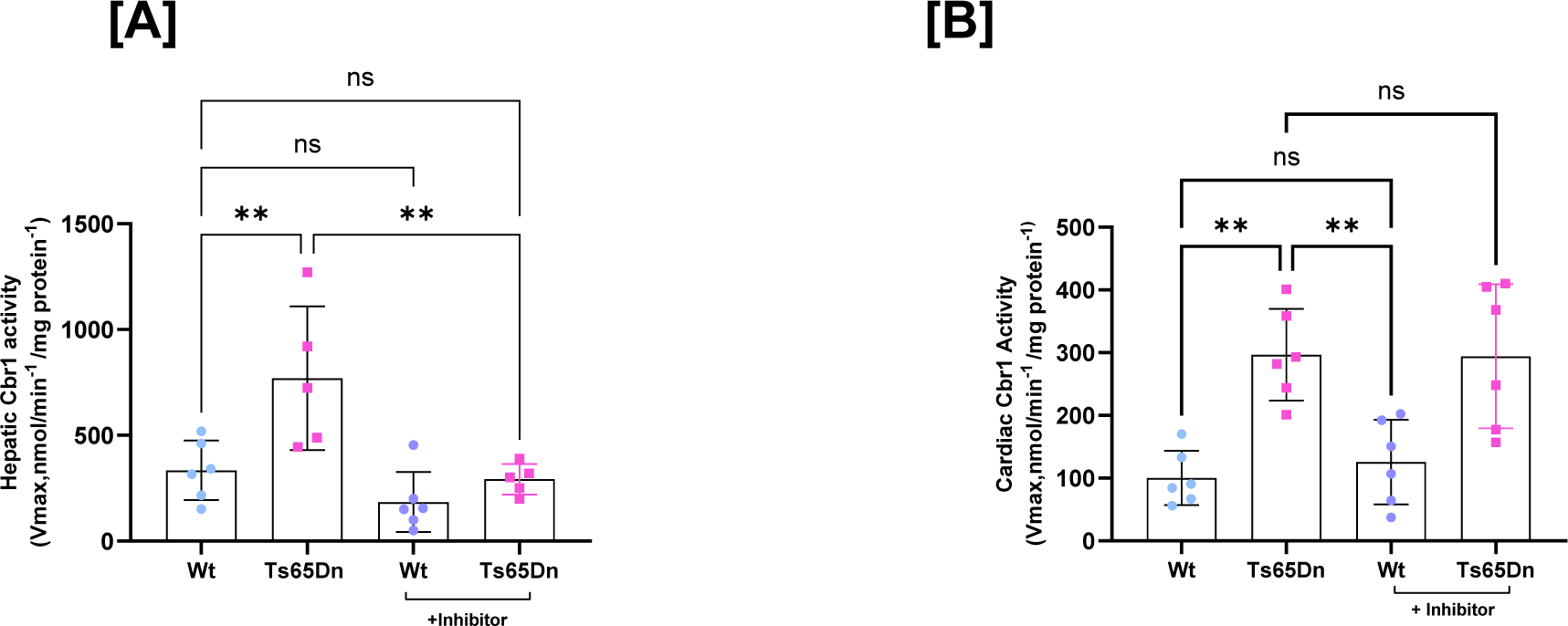
CBR1 activity as measured by doxorubicin reduction was increased in the liver and hearts of Ts65Dn mice compared to wildtype (Wt). Administration of hydroxy-PP-Me, an inhibitor of CBR1, reduced hepatic activity [A] in Ts65Dn mice but cardiac CBR1 activity was not affected [B]. Data are mean +/- standard deviation. (*P < 0.05, **P < 0.01, ***P < 0.001, and ****P < 0.0001)

Blood pressure was measured at baseline and during treatment with hydroxy-PP-Me. Median systolic, diastolic and mean arterial pressure during both the inactive period and active period were significantly lower in Ts65Dn mice compared with wild-type littermates (Table 2). Heart rate was significantly higher in the Ts65Dn mice compared with littermate controls (Table 2).

Cosinor analysis also showed that the MESOR (the rhythm adjusted-mean) of the systolic, diastolic and mean arterial pressure were significantly lower in Ts65Dn mice compared with wild-type littermates (Fig. 3, Table 3). Mesor of heart rate was significantly higher in the Ts65Dn mice compared with littermate controls (Fig. 3, Table 3). The amplitude of the circadian rhythm was not different between the groups for systolic pressure or heart rate. The amplitude of diastolic pressure and mean arterial pressure (MAP) was larger in the Ts65Dn mice compared with wild-type controls, corresponding to an increase in both active period blood pressure peak and inactive period blood pressure dip (Fig. 3, Table 3).

**Fig. 3:**
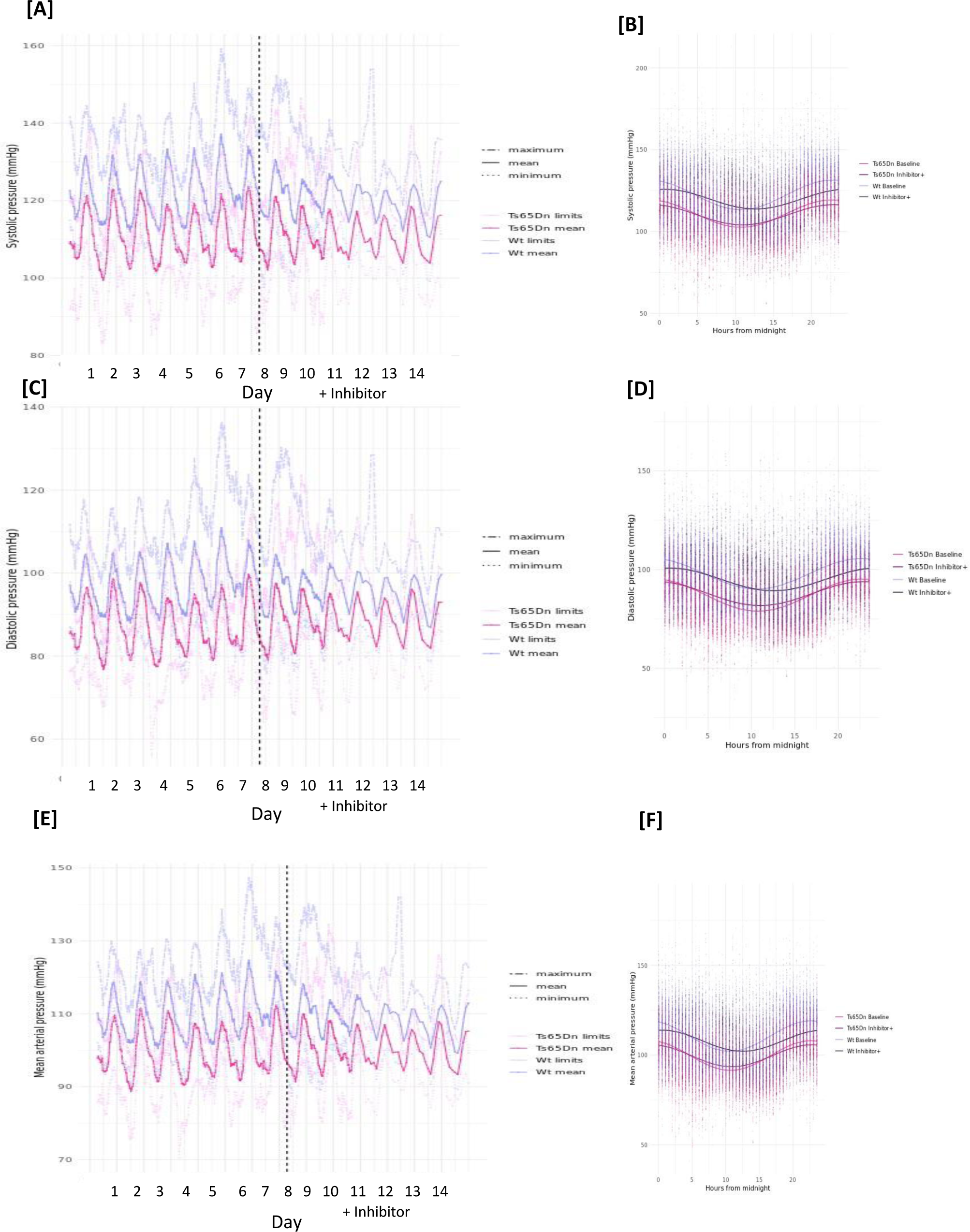
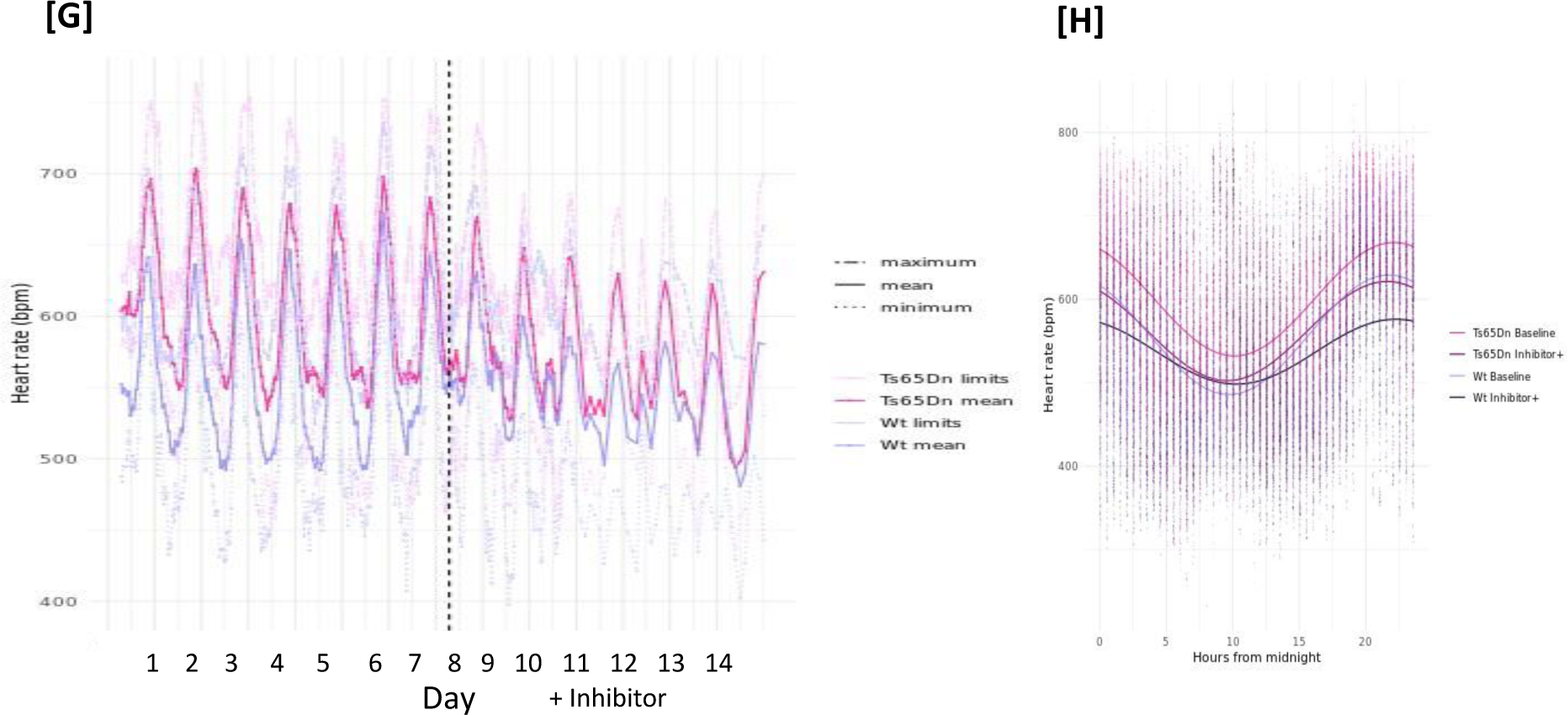
Ts65Dn mice have lower blood pressure and higher heart rate compared with Wt mice. [A, C, E and G] show five-hour rolling averages and minimum and maximum systolic, diastolic and mean arterial pressures, and heart rate measured by telemetry in Ts65Dn mice (pink) and their wild-type (Wt) littermate controls (purple) (n=8/group) over the course of 7 days of baseline measurements and then during daily treatment with CBR1 inhibitor hydroxy-PP-Me (+inhibitor) for 7 days. [B, D, F and H] show the cosinor curves fitted for the baseline and +inhibitor periods in Ts65Dn and Wt mice.

**Table 3:**
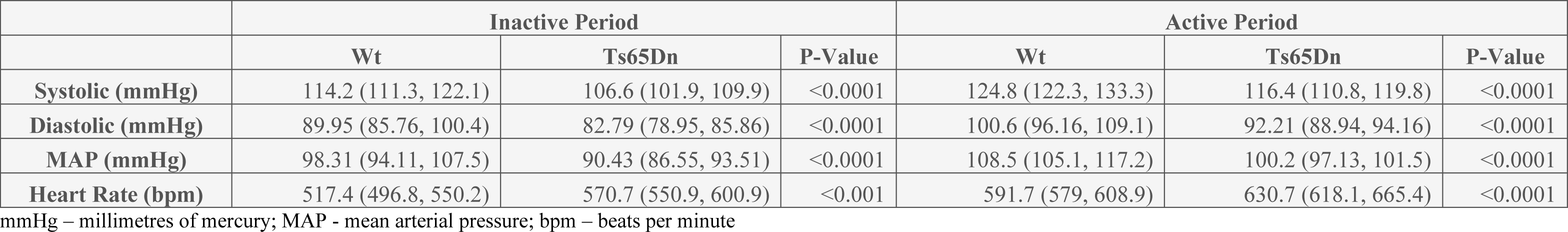
Blood pressure and heart rate of Ts65Dn mice and their wild-type littermate controls (Wt) during the inactive and active period. Data are median and interquartile range. Genotypes were compared using a Mann-Whitney U test.

Treatment with hydroxy-PP-Me, significantly increased the rhythm adjusted mean (MESOR) of systolic, diastolic and mean arterial pressure of Ts65Dn mice from baseline but decreased the MESOR in the wild type mice (Fig. 3, Table 4). There was a decrease in the amplitude of the rhythm in both wild-type and Ts65Dn mice corresponding to a reduction in the inactive phase dip in blood pressure i.e. inhibition of CBR1 blunted the fall in blood pressure (Table 4). Amplitude and MESOR of heart rate were significantly reduced by treatment in both groups of mice (Table 4).

**Table 4:** Cosinor analysis of blood pressure measured by telemetry in Ts65Dn mice and their wild-type littermate controls during the baseline period and during treatment with CBR1 inhibitor, hydroxy-PP-Me. The rhythm-adjusted mean (MESOR), amplitude of each parameter for each genotype during baseline and treatment and the outcome of statistical comparison are shown.

|  | Baseline |  |  | Treatment |  |  | Change with Treatment (P-Value) |  |
| --- | --- | --- | --- | --- | --- | --- | --- | --- |
|  | Wt | Ts65Dn | Wt vs Ts65Dn (P-Value) | Wt | Ts65Dn | Wt vs Ts65Dn (P-Value) | Wt | Ts65Dn |
| **SYSTOLIC** |  |  |  |  |  |  |  |  |
| MESOR (mmHg) | 122.09 | 110.87 | <0.001 | 120.01 | 111.42 | <0.001 | <0.001 | <0.01 |
| Amplitude (mmHg) | 8.97 | 8.84 | 0.13 | 5.90 | 6.17 | 0.19 | <0.001 | <0.001 |
| **DIASTOLIC** |  |  |  |  |  |  |  |  |
| MESOR (mmHg) | 97.47 | 87.07 | <0.001 | 95.26 | 87.99 | <0.001 | <0.001 | <0.001 |
| Amplitude (mmHg) | 7.68 | 8.53 | <0.001 | 5.61 | 6.14 | <0.01 | <0.001 | <0.001 |
| **MAP** |  |  |  |  |  |  |  |  |
| MESOR (mmHg) | 105.7 | 95 | <0.001 | 103.53 | 95.46 | <0.001 | <0.001 | <0.01 |
| Amplitude (mmHg) | 8.11 | 8.63 | <0.01 | 5.72 | 6.15 | <0.05 | <0.001 | <0.001 |
| **HEART RATE** |  |  |  |  |  |  |  |  |
| MESOR (bpm) | 557.5 | 602.5 | <0.001 | 539.8 | 564.2 | <0.001 | <0.001 | <0.001 |
| Amplitude (bpm) | 70.37 | 69.12 | 0.39 | 37.44 | 58.11 | <0.001 | <0.001 | <0.001 |

mmHg – millimetres of mercury; MAP – mean arterial pressure; bpm – beats per minute

### Mechanisms altering blood pressure

To determine if the blood pressure phenotype observed in *Cbr1^+/-^*mice was salt-sensitive, the animals were given a high-salt diet (3% sodium) and blood pressure was measured by telemtery for 7 days. During high salt feeding the mean systolic, diastolic and mean arterial pressure (MAP) increased in both groups but the difference between the groups remained constant (Table S1) demonstrating that salt sensitivity was similar between the groups. We confirmed that there were no differences in renal function as measured by glomerular filtration rate between *Cbr1^+/-^* mice and *Cbr1^+/+^* littermate controls (Fig. S2). Renal histology determined by light microscopy of haematoxylin and eosin-stained sections was normal in both genotypes (Fig. S2). The components of the renin-angiotensin-aldosterone system were not different between the groups (Fig. S3).

We then examined vascular function in *Cbr1^+/-^* animals and found no differences in the response of mesenteric vessels to vasoconstrictors or vasodilators to those of *Cbr1^+/+^* littermate controls (Fig. S4).

Plasma glucocorticoids (corticosterone and its inactive form 11-dehydrocorticosterone) measured by liquid chromatography tandem mass spectrometry were not different between the groups (Fig. S5).

Next, we examined known functions of CBR1 which may influence blood pressure by changing the vascular microenvironment. We explored the potential for CBR1 to impact oxidative stress, sympathetic tone and prostaglandin metabolism.

### Oxidative stress

CBR1 mediates detoxification of ROS making this a potential mechanism by which it influences blood pressure^45^. We therefore looked at measures of whole-body oxidative stress (total antioxidant capacity), lipid peroxidation (TBARS assay) and urinary 8-hydroxy-2’-deoxyguanosine (8-oxo-dG) as well as brain specific malondialdehyde (MDA). There were no differences in plasma or urinary measures of oxidative stress but brain MDA concentrations were increased in *Cbr1^+/-^*animals compared with *Cbr1^+/+^* littermate controls (Fig. 4).

**Fig. 4:**
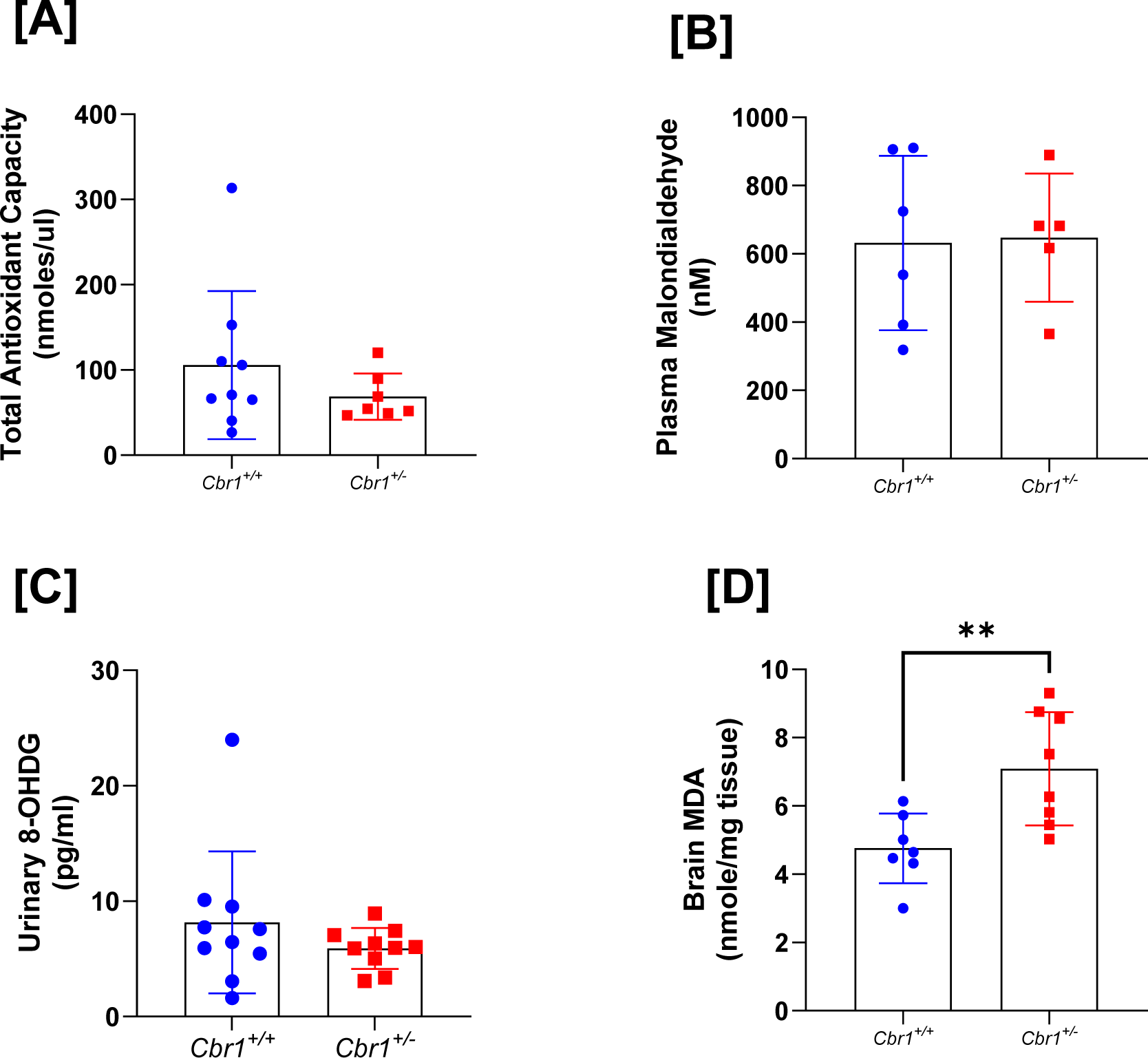
Oxidative stress markers were increased in the brains of mice heterozygous for Cbr1. Markers of oxidative stress, plasma malondialdehyde (MDA, TBARS assay) [A] and total antioxidant capacity and urinary 8-hydroxy-2’-deoxyguanosine (8-oxodG) [C] were not different between mice heterozygous for Cbr1 (Cbr1^+/-^) and their control littermates (Cbr1^+/+^) (n=8-11/group). Data are mean ± standard deviation (*P < 0.05, **P < 0.01, ***P < 0.001, and ****P < 0.0001)

### Sympathetic Activity

We determined if urinary excretion of catecholamines noradrenaline and adrenaline, as a proxy for sympathetic drive, were altered in *Cbr1^+/-^* compared with *Cbr1^+/+^* littermate controls. Urinary excretion of noradrenaline and adrenaline were measured in samples collected over a 24-hour period from mice housed in metabolic cages. Urinary excretion of noradrenaline but not adrenaline was increased in *Cbr1^+/-^* animals compared with their littermate controls (Fig. 5 A and C). We also showed that the mouse model of DS demonstrated decreased urinary excretion of noradrenaline but not adrenaline (Fig. 5 B and D). Administration of hydroxy-PP-Me normalised noradrenaline excretion in Ts65Dn animals (Fig. 5).

**Fig. 5:**
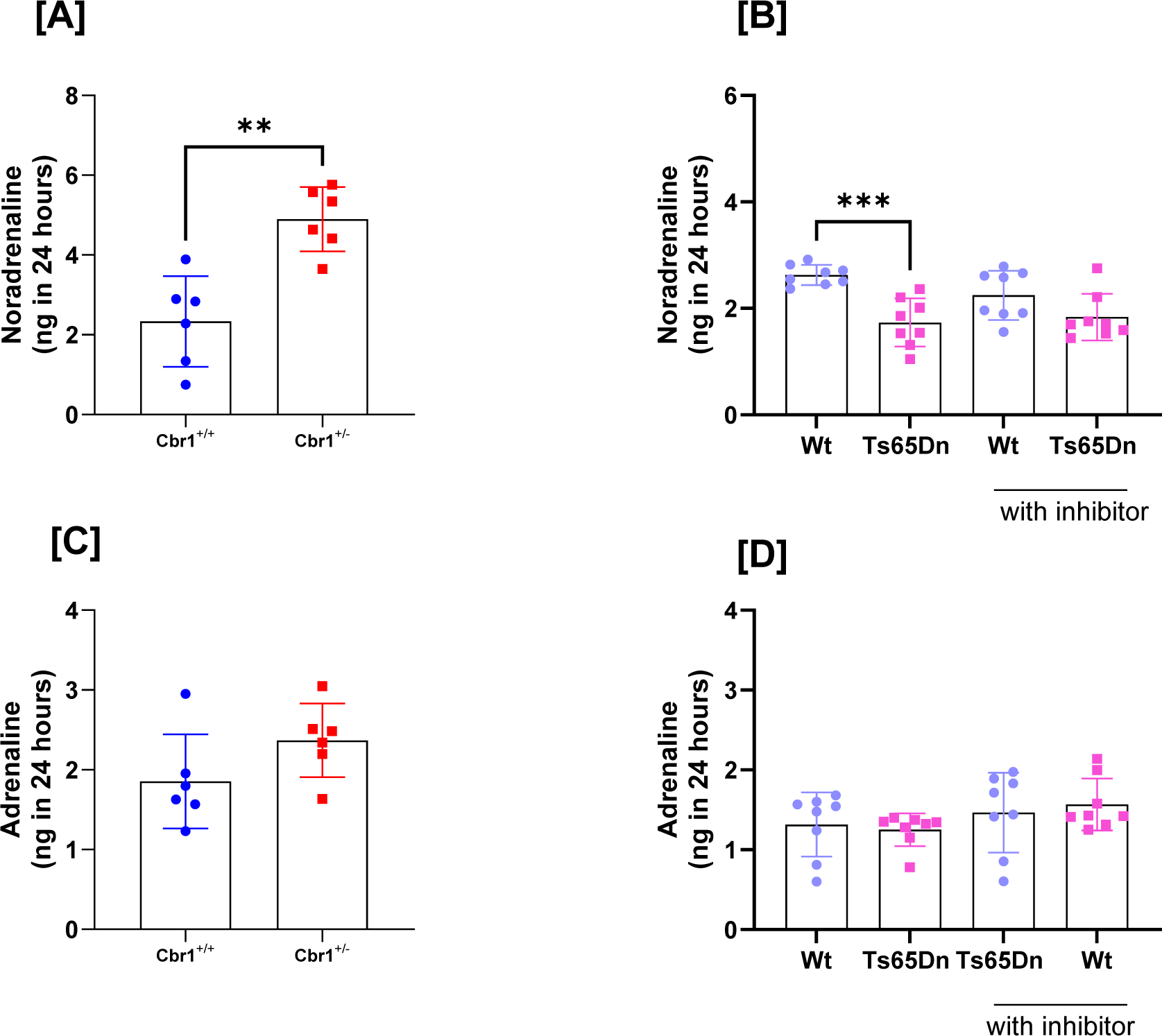
Cbr1 deletion results in increased sympathetic drive. Urinary noradrenaline excretion in a 24-hour period was increased in mice heterozygous for Cbr1 (Cbr1^+/-^) compared with their littermate controls (Cbr1^+/+^) [A] (n=6/group) and the opposite was true of Ts65Dn mice who had reduced noradrenaline excretion [B] (n=8/group). Urinary adrenaline excretion was not significantly different in Cbr1^+/-^ or Ts65Dn animals compared with wild-type controls [C, D]. Data are presented as group mean ± standard deviation (*P < 0.05, **P < 0.01, ***P < 0.001, and ****P < 0.0001).

### Prostaglandin metabolism

CBR1 inactivates prostaglandin E_2_ (PGE_2_) and converts it to prostaglandin F_2α_ (PGF_2α_), a mediator of blood pressure. As such we measured excretion of the substrate PGE_2_ in urine of mice heterozygous for *Cbr1* (*Cbr1*^+/-^) compared with their littermate controls (*Cbr1*^+/+^) and found that *Cbr1*^+/-^ mice had increased excretion indicating reduced systemic metabolism (Fig. 6). The opposite was true of Ts65Dn animals compared with littermate controls, but this was normalised by administration of hydroxy-PP-Me (Fig. 6).

**Fig. 6:**
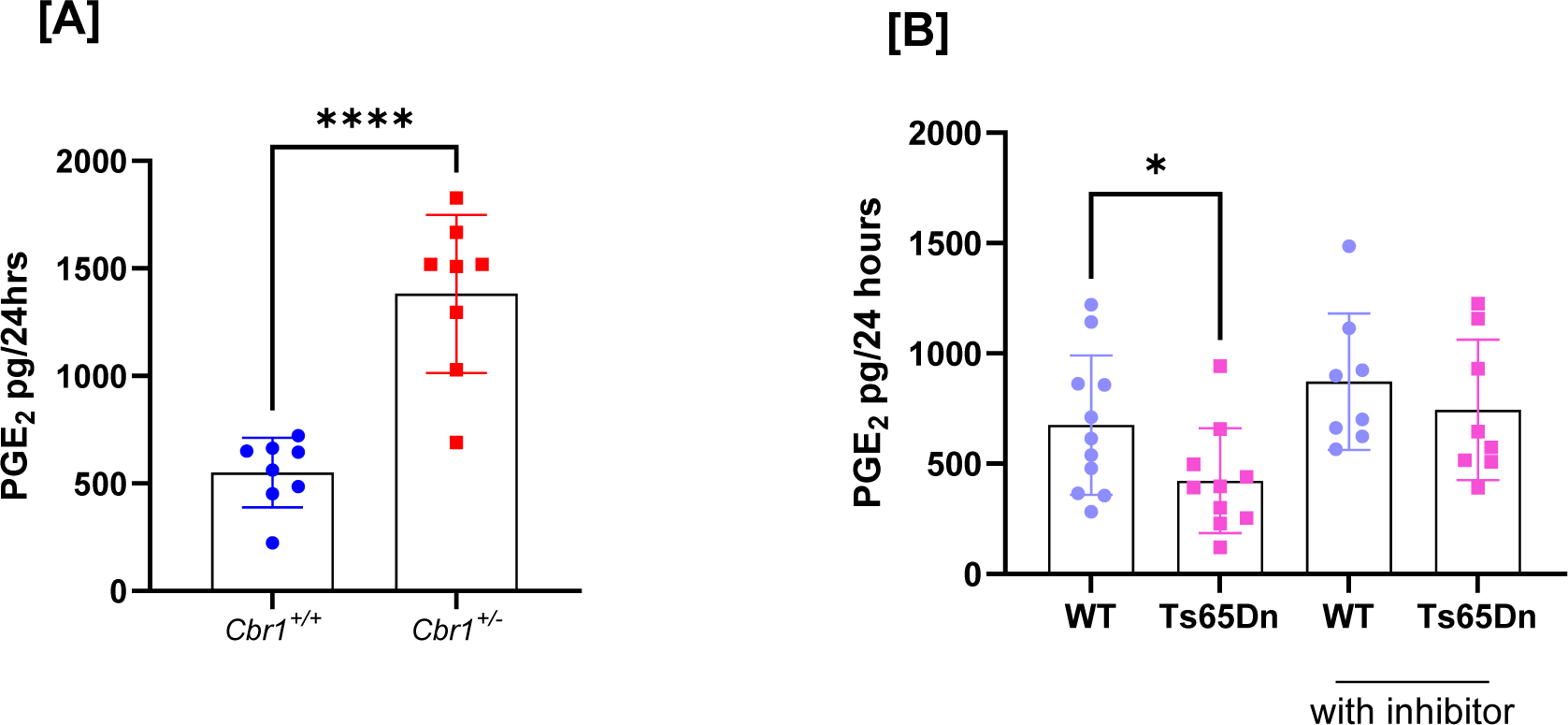
Prostaglandin metabolism is altered by Cbr1 deletion and by CBR1 inhibition. [A] Urinary prostaglandin E_2_ excretion was increased in mice heterozygous for Cbr1 (Cbr1^+/-^) compared with littermate controls (Cbr1^+/+^) (n=8/group). PGE_2_ excretion was decreased in Ts65Dn animals compared to controls and this was corrected by administration of the inhibitor[B] (n=8-11/group). Data are presented as group mean ± standard deviation (*P < 0.05, **P < 0.01, ***P < 0.001, and ****P < 0.0001).

## Discussion

In this study we present the first demonstration of carbonyl reductase 1 (CBR1) as a novel regulator of blood pressure. Our data indicate that increased CBR1 contributes to hypotension observed in a mouse model of Down Syndrome. Mice heterozygous for *Cbr1,* with a 50 % reduction in enzyme activity in all tissues^28^, had increased systolic, diastolic and mean arterial pressure. In the absence of changes in renal function, salt sensitivity or vascular reactivity, the most plausible drivers of altered blood pressure are the observed alterations in sympathetic tone and prostanoid metabolism, inferred from urinary catecholamine and prostaglandin excretion.

PGE_2,_ a substrate of CBR1^46,47^, is known to induce hypertension and catecholamine release when administered intracerebroventricularly to rats^48^ and yet have the opposite effect when given systemically^49^ where it also increases renin secretion^50^. Reduced levels of PGE_2_ in the brain are found in the Ts1Cje rodent model of Down Syndrome and this is reversed when the copy number of the *Cbr1* gene is restored^51^. Our results are consistent with this, demonstrating that mice with reduced CBR1 activity had reduced metabolism (and hence increased excretion) of PGE_2_. We did not identify the source of this increased PGE_2_ but given we did not see differences in plasma renin, and systemic vascular function was unaffected, we might suppose that the increases were localised, possibly in the brain and therefore possibly influencing sympathetic activity.

CBR1 is known to reduce oxidative stress centrally^52^ and this was apparent in our study which showed increased brain oxidative stress in mice deficient in *Cbr1*. Interestingly we found no evidence of a systemic increase in markers of oxidative stress in *Cbr1^+/-^*mice which is consistent with the normal vascular and renal function we saw in these animals. It is also likely that compensatory mechanisms come into play when *Cbr1* is lacking or that 50 % of normal levels are sufficient to protect cells elsewhere.

It is suggested that, in DS, blunted sympathetic control results in a reduced response to exercise associated with exercise intolerance and low VO_2_ max^17,53^ and is also implicated in sleep apnoea in these patients^11^. Our study suggests that activity of CBR1 impacts sympathetic tone, particularly noradrenaline release. In addition to indirect effects via oxidative stress or prostaglandin metabolism *Cbr1* could also directly affect the sympathetic nervous system; for example, it was recently described as the predominant pathway by which the endogenous monoamine oxidase inhibitor, isatin, is inactivated^24,54^.

Hypotension can have a significant impact on quality of life for people with DS, limiting exercise, contributing to sleep disturbances and potentially accelerating the onset and progression of Alzheimer’s disease. There are currently no specific treatments available as the pathophysiology is unknown. Despite inhibition of CBR1 with hydroxy-PP-Me resulting in tissue-specific enzyme inhibition there was still a blunting of the normal inactive phase dip in blood pressure and an increase in noradrenaline excretion and prostaglandin metabolism in this mouse model of DS. This suggests there is merit in pursuing CBR1 inhibition by this or other compounds^55,56^ as a therapeutic intervention in patients for whom hypotension impacts quality of life. It is interesting to note that inhibition of CBR1 reduced blood pressure in the wild type mice in whom CBR1 levels were “normal” so it seems likely that a critical balance of CBR1 activity is required to maintain a normal vascular microenvironment and blood pressure; as such, partial inhibition may be an attractive therapeutic option.

Whilst we have focused on the role of *Cbr1* in DS, our work has wider implications. In the general population there is wide variation in CBR1 expression and activity levels between the sexes and between ethnic groups^57^ and our data suggest that *Cbr1* may be a novel gene influencing blood pressure. Inhibitors of CBR1, particularly flavonoids, exist in many foodstuffs and food supplements^58^ and are often advocated as supplements for people with metabolic disease. Pharmacological inhibitors of CBR1 are being explored for use as adjunctive therapy in chemotherapeutic regimes which include doxorubicin because CBR1 metabolises doxorubicin to cardiotoxic daunorubicin which limits its use, particularly in DS patients^21,35,40^. Our data suggest that inhibition of *Cbr1* should be used with caution in those with or susceptible to hypertension.

This study has identified a role for CBR1 in the control of blood pressure and in sympathetic drive. Down syndrome is an incredibly complex disorder and there is a well-described dose dependent effect of the trisomy but our data suggest that CBR1 may be a potential therapeutic target in those DS patient for whom low blood pressure impacts their quality of life.

### Limitations

It is important to acknowledge the limitations of these studies. We used mice which were heterozygous for *Cbr1* in every tissue, we therefore cannot ascertain which tissue is most important in the hypotensive phenotype. We acknowledge the limitations of inferences made in mice in such a complex human syndrome as DS, the role or importance of Cbr1 in human blood pressure control may differ from that in mice.

## List of Supplementary Materials

Materials and Methods

Table S1 to S4

Fig. S1 to S5

References 59 and 60

## Acknowledgements

The authors would like to acknowledge the staff of the University of Edinburgh animal facility.

## Sources of Funding

This work and author RM were funded by a Wellcome Trust Clinical Research and Career Development Fellowship (206587/Z/17/Z). Author BRW was supported by a a Wellcome Senior Investigator Award.

## Disclosures

None

## Notes

### Competing Interest Statement

The authors have declared no competing interest.

